# *Mkk4* and *Mkk7* are important for retinal development and axonal injury-induced retinal ganglion cell death

**DOI:** 10.1101/292482

**Authors:** Rebecca L. Rausch, Stephanie B. Syc-Mazurek, Kimberly A. Fernandes, Michael P. Wilson, Richard T. Libby

## Abstract

The mitogen activated protein kinase (MAPK) pathway has been shown to be involved in both neurodevelopment and neurodegeneration. c-Jun N-terminal kinase (JNK), a MAPK shown to be important in retinal development and after optic nerve crush injury, is regulated by two upstream kinases: MKK4 and MKK7. The specific requirements of MKK4 and MKK7 in retinal development and retinal ganglion cell (RGC) death after axonal injury, however, are currently undefined. Optic nerve injury is an important insult in many neurologic conditions including traumatic, ischemic, inflammatory, and glaucomatous optic neuropathies. Mice deficient in *Mkk4*, *Mkk7*, and both *Mkk4* and *Mkk7* were generated. Immunohistochemistry was used to study the distribution and structure of retinal cell types and to assess RGC survival after optic nerve injury (mechanical controlled optic nerve crush; CONC). Adult *Mkk4* and *Mkk7* deficient retinas had all retinal cell types. With the exception of small areas of lamination defects with photoreceptors in *Mkk4* deficient mice, the retinas of both mutants were grossly normal. Deficiency of *Mkk4* or *Mkk7* reduced JNK signaling after axonal injury in RGCs. *Mkk4* and *Mkk7* deficient retinas had a significantly greater percentage of surviving RGCs 35 days after CONC as compared to wildtype controls (*Mkk4*: 51.5%, *Mkk7*: 29.1% WT: 15.2%; p<0.001). Combined deficiency of *Mkk4* and *Mkk7* caused failure of optic nerve formation, irregular retinal axonal trajectories, disruption of retinal lamination, clumping of RGC cell bodies, and dendritic fasciculation of dopaminergic amacrine cells. These results suggest that MKK4 and MKK7 may serve redundant and unique roles in molecular signaling important for retinal development and injury response following axonal insult.

## Introduction

The mitogen activated protein kinase (MAPK) pathway is involved in development, neurodegeneration, and the immune response^1,2,3,4,5^. In the retina, MAPK signaling plays a role in retinal formation and axonal injury-induced retinal ganglion cell (RGC) death^6,7,8,9,10,11,12^. Classical MAPK signaling consists of a three-tiered cascade, in which sequential phosphorylation by upstream MAP kinases (MAP3Ks, MAP2Ks, and MAPKs) leads to differential cellular responses at the transcriptional level^3,13^. c-Jun N-terminal kinase (JNK) is regulated by two upstream MAP2Ks: MKK4 and MKK7^5,13,14^. However, the specific requirements of MKK4 and MKK7 in retinal development and neurodegeneration are currently undefined.

MKK4 and MKK7 are required for normal development^15^. In the central nervous system, MKK4 and MKK7 and their downstream effector molecules, the JNKs (JNK1, 2 and 3), have been shown to play important roles in both development and maintenance of neural structures. MKK4, MKK7, and the JNKs contribute to the regulation of cellular organization and axonal migration^16,17,18^. Such contributions have been suggested to occur through both overlapping and non-redundant mechanisms^19^. JNK signaling has been shown to contribute to multiple aspects of retinogenesis such as progenitor cell proliferation^14,20^. The exact contributions of MKK4 and MKK7 to retinal development, however, remain largely unexplored. In the adult eye, MAPK signaling is critical for the apoptotic injury response in RGCs after axonal injury. Both upstream (MAP3K) and downstream MAPK members have been shown to be key mediators of RGC death after axonal injury. Specifically, JNKs and their canonical downstream effector molecule, the transcription factor JUN, are important for RGC death after mechanical- and ocular hypertension-induced axonal injury^7,8,21,22,23,24^. Despite this known involvement, the critical molecular events leading from axonal injury to RGC death are not fully defined. Determining the molecular mechanisms of RGC pro-death signaling after axonal injury is necessary for understanding the molecular underpinnings of diseases such as glaucoma and traumatic optic neuropathies which result in RGC loss.

The importance of JNK signaling for both RGC development and response to axonal injury is well established, however little is known regarding the role of the MAP2Ks upstream of JNK in these processes. Selectively targeting these upstream MAPKs may allow us to define the specific pathological signaling pathway that activates pro-death JNK activation in RGC axons after an insult. Furthermore, understanding the contribution of MKK4 and MKK7 to the injury response and to JUN activation in RGCs will likely have implications for other diseases or traumas involving axonal injury. Here, using conditional null alleles of *Mkk4* and *Mkk7*, we investigate the importance of these two MAP2Ks for RGC maturation and response to axonal injury. We demonstrate these MAP2Ks have redundant and unique roles in molecular signaling important for retinal development and injury signaling.

## Materials and Methods

### Mice

All experiments were conducted in adherence with the Association for Research in Vision and Ophthalmology’s statement on the use of animals in ophthalmic and vision research and were approved by the University of Rochester’s University Committee on Animal Resources. Animals were housed on a 12-hour light and dark cycle and received chow and water ad libitum. Floxed alleles of *Mkk4* and *Mkk7*^25,26^ were maintained on the C57BL/6J background. To generate animals deficient in *Mkk4* or *Mkk7*, mice carrying the respective floxed allele were bred to animals carrying *Six3*cre which is first expressed in the optic cup between E9.0-9.5^27^. Animals carrying the Cre recombinase and one *Mkk4* or *Mkk7* floxed allele were intercrossed to generate animals: 1) carrying Cre recombinase and two copies of either *Mkk4* or *Mkk7* floxed alleles, referred to as *Mkk4* deficient (*Mkk4*^*-/-*^ or *Six3*cre^+^*;Mkk4*^*f/f*^) and *Mkk7* deficient (*Mkk7*^*-/-*^ or *Six3*cre^+^;*Mkk7*^*f/f*^) respectively and, 2) animals without floxed alleles and/or without Cre recombinase, referred to as wildtype controls (*Six3*cre^*+*^;*Mkk4*^*+/+*^, *Six3*cre−;*Mkk4*^*+/+*^, *Six3*cre−;*Mkk4*^f/+^, *Six3*cre−;*Mkk4*^*f/f*^, *Six3*cre^*+*^, *Mkk7*^*+/+*^, *Six3*cre−, *Mkk7*^*+/+*^, *Six3*cre−;*Mkk7*^*f/+*^, or *Six3*cre−;*Mkk7*^*f/f*^). Animals deficient in both *Mkk4* and *Mkk7* where generated by breeding animals carrying the floxed alleles and *Six3*cre together to generate *Six3*cre^*+*^*;Mkk4*^*f/f*^;*Mkk7* ^*f/f*^ animals (*Mkk4/Mkk7* deficient animals).

### Retinal Histology, Immunohistochemistry, and Cell Counts

Eyes used to evaluate retinal morphology were processed in 2.5% glutaraldehyde; 2% paraformaldehyde (PFA) for 24 hours at 4°C. Eyes were then dehydrated with a series of ethanol washes, embedded in Technovit (Electron Microscopy Services), sectioned at 2.5μm, and stained with Multiple Stain Solution (Polysciences, Inc). Eyes to be processed for immunohistochemistry and retinal flat mounts were harvested and processed in 4% PFA for two hours at room temperature prior to storage in 1M PBS (phosphate buffered saline). As has been previously described, the anterior segment of the eye was dissected away and the posterior segment of the eye was processed for cyrosectioning (14μm sections) or whole retina flat mounts ^7^. To evaluate gross morphology of the optic nerve, 8 additional animals per genotype were perfused with 4% PFA. Brains were then dissected from the skull and the ventral surface containing both optic nerves was photographed using a stereomicroscope.

For immunohistochemical staining on adult retinas, cryosections were blocked in 10% horse serum with 0.1% TritonX in 1xPBS for 3 hours at room temperature and then incubated in primary antibody overnight at 4°C. Primary antibodies included: goat anti SOX2 (1:250, Santa Cruz), goat anti CHaT (1:200, Millipore), rabbit anti Calretinin (1:1000, Celco), mouse anti Calbindin-D-28K (1:1000, Sigma), rabbit anti PKCα (1:2000, Sigma), and mouse anti pJNK (1:250, Cell Signaling). Cryosections were washed in PBS and incubated in Alexafluor-conjugated secondary antibodies (Invitrogen), for 2 hours at room temperature before being washed in PBS. Cyrosections were then counterstained with DAPI and mounted in Fluorogel in TRIS buffer (Electron Microscopy Sciences).

For immunohistochemistry on flat mounts, floating retinas were blocked in 10% horse serum with 0.3% TritonX in 1xPBS overnight on a shaker at 4°C and then incubated in primary antibody for three days at 4°C as previously described^28^. Primary antibodies included: mouse anti TUJ1 (1:1000, BioLegend), goat anti BRN3b (1:250, Santa Cruz), rabbit anti TH (1:1000 Millipore), rabbit anti pJUN (1:250, Cell Signaling), rabbit anti cCASP3 (1:1000, Millipore), and rabbit anti Neurofilament light chain (1:100; Cell Signaling). Whole retinas were then washed in 1XPBS and incubated in Alexafluor-conjugated secondary antibodies for two days at room temperature. Whole retinas were then mounted RGC side up in Fluorogel in TRIS buffer. RGCs were quantified in eight equally spaced 40x fields taken approximately 220μm from the peripheral edge of the retina as previously described^29^. Quantification was completed using the cell counter tool in ImageJ.

### Protein Extraction and Western Blotting

Western blots were performed as previously described ^8^. Briefly, retinas were dissected and placed in 100 µl ice cold lysis buffer (1X RIPA buffer (Santa Cruz, 24948) and protease/phosphatase inhibitor cocktail (Cell Signaling, 5872S)). Tissue was lysed by sonication (Bransa Digital Sonifier, 10% amplitude for 3 seconds) prior to spinning down cellular debris in a microcentrifuge (10,000 RPM, 4°C, 5 minutes). 10 µl of supernatant was combined with 10 µl 2X Laemmli loading buffer (Bio-Rad) and boiled for 10 minutes and run on a 12% SDS-PAGE gel. After transfer to a PVDF membrane, membranes were treated with a Qentix Western Blot Signal Enhancer kit (Thermo Scientific, 21050) according to manufacturer’s instructions. Membranes were blocked and probed overnight at 4°C with one of the following primary antibodies: rabbit anti-pJUN (1:250, Cell Signaling), rabbit anti-pJNK (1:500, Cell Signaling), rabbit anti-MKK4 (1:500, Cell Signaling), rabbit anti-MKK7 (1:500, Cell Signaling), or rabbit anti-GAPDH (1:2000, Cell Signaling). The following day, membranes were washed and probed with secondary antibody: HRP-conjugated anti-rabbit IgG (1:5000, Bio-Rad). Immunoreactive bands were detected using a chemiluminescence kit (Immun-star, Bio-Rad 170-5070) prior to exposure using either film or digital detection equipment (Azure Biosystems c500). Membranes were occasionally stripped following development and treated with another primary antibody (stripping buffer: 0.1 M Tris-Cl pH 6.8, 2% SDS, 0.7% β-mercaptoethanol). Densitometric analysis was conducted using ImageJ software, and pixel densities of experimental bands were normalized to those of GAPDH loading controls.

### Mechanical Optic Nerve Injury

Controlled optic nerve crush (CONC) was performed as previously described^28,30^. Prior to the procedure, animals were anesthetized with 100mg/kg ketamine and 10mg/kg xylazine. Briefly, the optic nerve was surgically exposed and then crushed with a pair of self-closing forceps immediately behind the eye for 5 seconds. A cohort of eyes underwent sham surgery, in which the optic nerve was exposed but not crushed. These eyes along with eyes that had not been manipulated served as experimental controls. Following CONC or sham surgery, animals were harvested after 2 hours, 1 day, 5 days, or 35 days.

### Statistical Analysis

At least three retinas of each genotype were analyzed for all experimental conditions. Experimenters were masked to genotype and experimental condition for all quantification of RGC counts. Unpaired Student’s t-tests were used to compare differences across two groups. A one-way ANOVA followed by the Bonferroni *post hoc* test for group comparisons was used to compare differences across more than two groups at a single time point. A *P* value <0.05 was considered statistically significant. Means +/-SEM are displayed in graphs.

## Results

### Deficiency of *Mkk4* or *Mkk7* leads to mild alterations in retinal structure

To generate retinas deficient in *Mkk4* or *Mkk7*, *Six3*cre (a retina-specific cre) was used recombine floxed alleles of either *Mkk4* or *Mkk7*^27^. Western blot analysis demonstrated efficient deletion of both *Mkk4* and *Mkk7* with >95% and >85% protein reduction, respectively (Fig. S1). Previously, other members of the MAPK signaling family, including JNK, have been shown to be necessary for retinal development^12,31^. To determine if deletion of *Mkk4* or *Mkk7* is necessary for retinal development, semi-thin sections of adult WT, *Mkk4* deficient, and *Mkk7* deficient retinas containing the optic nerve head were evaluated (Fig. 1A). The *Mkk4* deficient retina and optic nerve head had grossly normal morphology, with the exception of occasional photoreceptor cell bodies misplaced amongst photoreceptor inner and outer segments. Also, there appeared to be more cells in the inner plexiform layer in the *Mkk4* deficient retinas than in control retinas. The *Mkk7* deficient retinas had increased areas of cellularity in the surface nerve fiber layer and prelaminar region of the optic nerve head. Retinal lamination in *Mkk7* deficient animals appeared normal, although there was slight disruption in the contour of the inner limiting membrane and ganglion cell layer.

**Figure 1.**
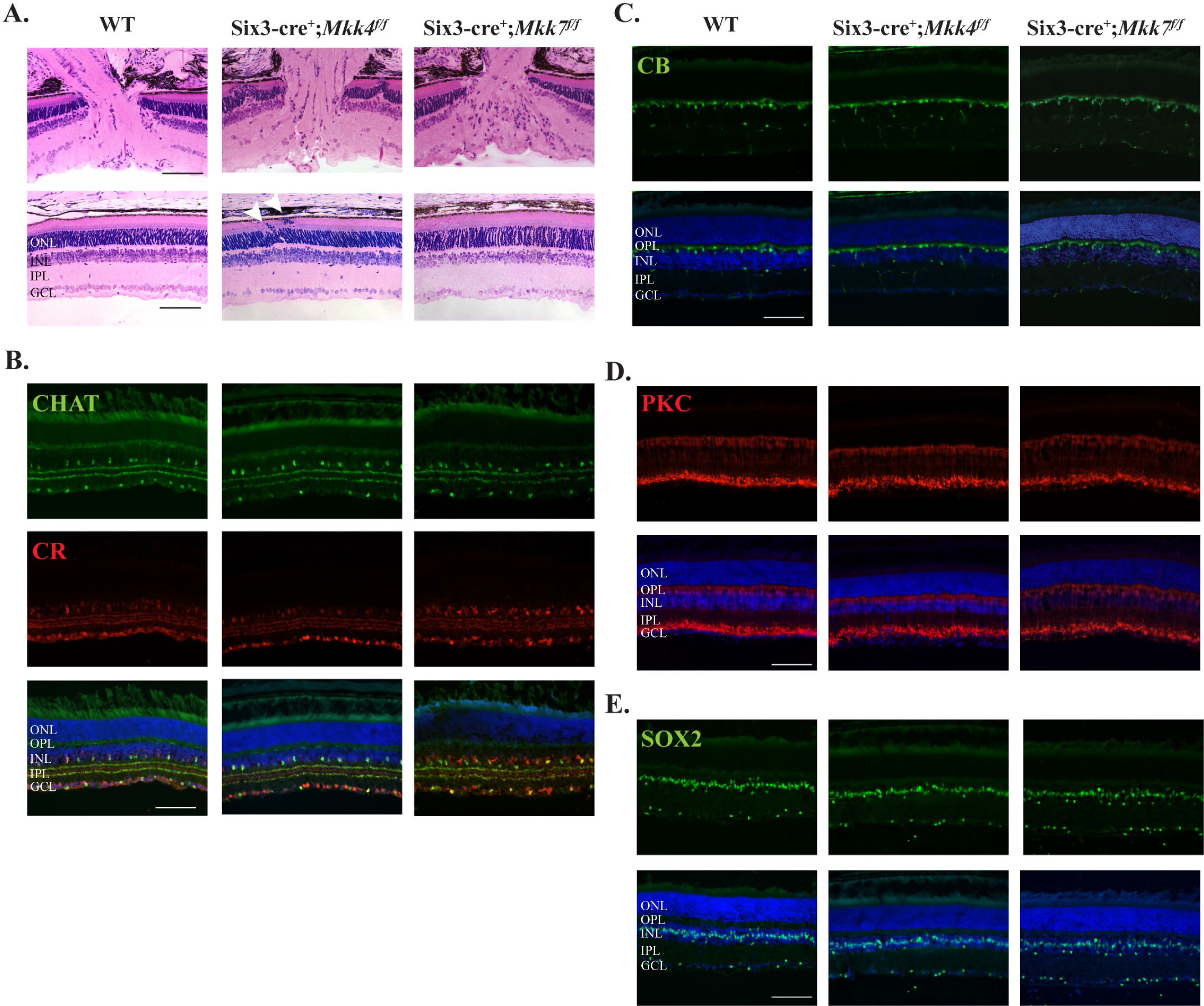
Adult *Mkk4* and *Mkk7* deficient retinas exhibit presence of all major cell types with minor structural alterations. (**A**) Representative 3μM plastic sections stained with H&E demonstrate areas of photoreceptor cell body attachment to retinal pigmented epithelium in *Mkk4* deficient retinas (arrowheads). Misplaced cells are also seen in the inner plexiform layer in *Mkk4* deficient retinas. Retinal lamination in *Mkk7* deficient retinas appeared grossly normal, though regions of the optic nerve head displayed slightly increased cellularity in the prelaminar region. Antibodies against ChAT and calretinin revealed normal development of cholinergic and AII amacrine cells, respectively, in both *Mkk4* and *Mkk7* deficient retinas (**B**). Calbindin-D-28K labeled horizontal cells (**C**), PKCα labeled bipolar cells (**D**), and SOX2 labeled Müller glia (**E**) in *Mkk4* and *Mkk7* mutants were furthermore indistinguishable from controls. N ≥ 4 retinas for each genotype. Scale bars: 100μm.

To further examine if *Mkk4* and *Mkk7* were necessary for retinal development, immunohistochemistry was used to study specific retinal cell types. Antibodies against choline acetyltransferae (ChAT) and calretinin were used to label amacrine cell bodies in the inner nuclear layer and synaptic strata in the inner plexiform layer. ChAT labels cholinergic amacrine cells and synaptic layers 2 and 4 whereas calretinin labels AII amacrine cells and layers 2, 3, and 4 in the inner plexiform layer^32^. Amacrine cells and inner plexiform lamination in both *Mkk4* and *Mkk7* deficient retinas appeared to develop normally (Fig. 1B). Horizontal cells and bipolar cells, stained with calbindin D-28K and PKCα, respectively, also appeared normal (Fig. 1C, D). Finally, there were no observable differences between WT and *Mkk4* or *Mkk7* deficient retinas in SOX2 staining, which labels Müller glia and a subset of amacrine cells (Fig. 1E). Together, these data suggest that aside from sporadic photoreceptor nuclei directly abutting retinal pigment epithelium in the outer nuclear layer, all major cell types within the inner nuclear layer appeared to grossly differentiate properly.

### Adult *Mkk4* and *Mkk7* deficient animals have fewer RGCs

In the semi-thin sections (Fig. 1A), the cellularity of the ganglion cell layer appeared less in the *Mkk4* and *Mkk7* deficient retinas than in the controls. There also appeared to be some disorganization in the nerve fiber layer or the inner limiting membrane in the *Mkk7* deficient retina. Therefore, the number of TUJ1+ RGCs was counted in retinal flat mounts in wildtype, *Mkk4* deficient, and *Mkk7* deficient retinas (Fig. 2A). Retinas contained 15% and 25% fewer RGCs in *Mkk4* and *Mkk7* deficient animals, respectively, as compared to their wildtype controls (P<0.05 for each comparison). Furthermore, while no gross alteration was observed in *Mkk4* deficient RGCs, *Mkk7* deficiency resulted in sporadic clumping and axonal fasciculation (discussed below; experimenters performing cell counts avoided these small areas). In order to determine whether the decreased RGC density in adult *Mkk4* and *Mkk7* deficient retinas was due to an early developmental defect, flat mounts were examined at P0, a time point subsequent to RGC birth and determination ^33^. Retinas were stained for BRN3B (POU4F2), another marker for RCGs^34,35^. RGC cell counts in both mutant mice were normal at this age, suggesting the correct amount of RGCs were born but died at a later stage (Fig. 2B).

**Figure 2.**
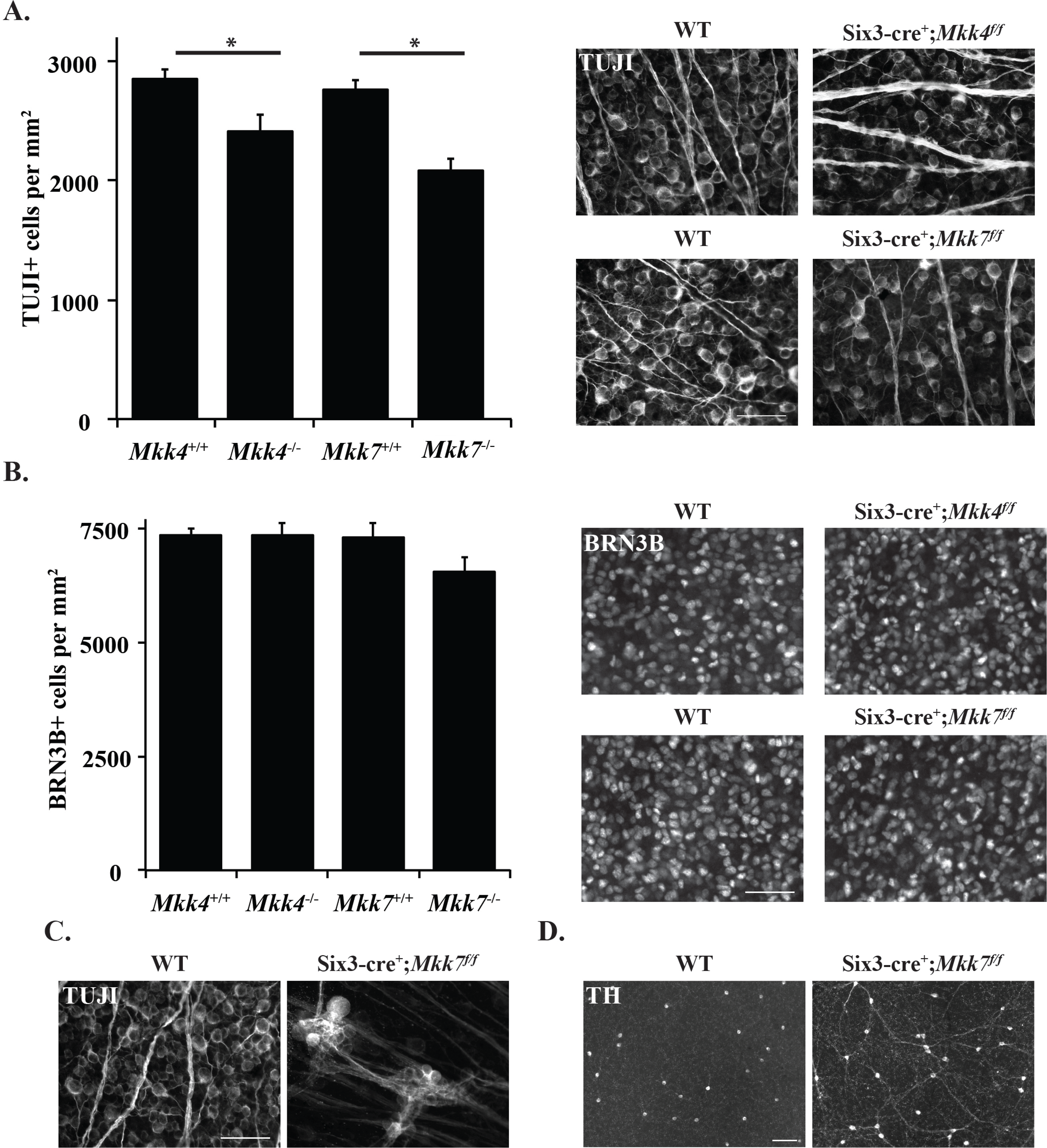
Adult *Mkk4* and *Mkk7* deficient animals have fewer retinal ganglion cells resulting from postnatal dropout. (**A**) The number of TUJ1 positive RGCs is significantly reduced in adult *Mkk4* and *Mkk7* deficient retinal flat mounts. Representative flat mounts are displayed to the right of the graph. (*, p ≤ 0.05). N ≥10 retinas per genotype. (**B**) P0 flat mounts stained with BRN3B reveal normal RGC densities in both *Mkk4* and *Mkk7* deficient retinas, implying RGC dropout occurs postnatally. Representative flat mounts shown to the right of the graph. N≥4 retinas per genotype. All examined *Mkk7* deficient adult retinas displayed areas of abnormal fasciculation in both TUJ1 positive RGCs (**C**; N≥12) and tyrosine hydroxylase (TH) positive amacrine cells (**D**; N≥4). Scale bars: (**A-C**), 50μm; (D), 100μm. Error bars represent S.E.M.

### Deficiency of *Mkk4* or *Mkk7* causes axonal fasciculation in RGCs and some amacrine cells

RGC cell body clumping and axonal fasciculation was observed in 100% of *Mkk7* deficient retinas examined and never observed in *Mkk4* deficient retinas (Fig. 2C). Areas of affected retina were judged to account for only a small portion of the ganglion cell layer (less than 1%). Aside from the few areas of clumping and fasciculation, RGC structure in *Mkk7* deficient retinas appeared similar to WT controls. JNK1, a downstream target of MKK4 and MKK7, is required for Netrin-1-induced axon outgrowth in the spinal cord^36^. Down syndrome cell adhesion molecule (DSCAM) has also been implicated in JNK1-Netrin-1 mediated neurite growth, and its loss of function leads to clustering of both RGC axons as well as cholinergic amacrine cell dendrites^36,37^. To examine whether the same amacrine cell phenotype resulted from *Mkk4* or *Mkk7* deficiency, additional retinal flat mounts were stained for tyrosine hydroxylase (TH), marking a subset of dopaminergic amacrine cells. Similar to what was observed in *Dscam* mutant mice, deficiency of *Mkk7* resulted in fasciculation of dopaminergic amacrine cell dendrites (Fig. 2D). Dopaminergic cell fasciculation was far less pronounced within *Mkk4* deficient retinas (data not shown).

### Deficiency of *Mkk4* or *Mkk7* does not prevent JNK-JUN signaling after axonal injury in RGCs

MKK4 and MKK7 are the only two MAP2Ks that activate JNK, which in turn activates its conical target, JUN. JNK-JUN signaling is important for pro-apoptotic signaling after axonal injury^6,7,38,39,40,41^. MKK4 and MKK7 have been previously reported to serve non-redundant functions *in vivo*^18,26,42,43,44^. For example, activation of MKK4 but not MKK7 triggers neuronal death following oxidative stress^45^ while MKK7 is required for JNK activation caused by pro-inflammatory cytokines (such as TNFα and IL-1)^15^. Together these results suggest that activation of JNK in the setting of axonal injury may be preferentially controlled by a single MAP2K. To test this possibility, JNK-JUN signaling was evaluated in the *Mkk4* and *Mkk7* deficient animals. Activated JNK (pJNK) and JUN (pJUN) are upregulated following optic nerve injury^7,10,46,47^. In retinal sections, expression of pJNK was observed in RGC axons entering the optic nerve head in WT, as well as in *Mkk4* and *Mkk7* deficient mice (Fig. 3A). Similarly, pJUN positive RGCs were observed in WT, *Mkk4* and *Mkk7* deficient flat mounts (Fig. 3B). Thus, neither *Mkk4* nor *Mkk7* was independently necessary for activation of JNK or JUN after axonal injury.

**Figure 3.**
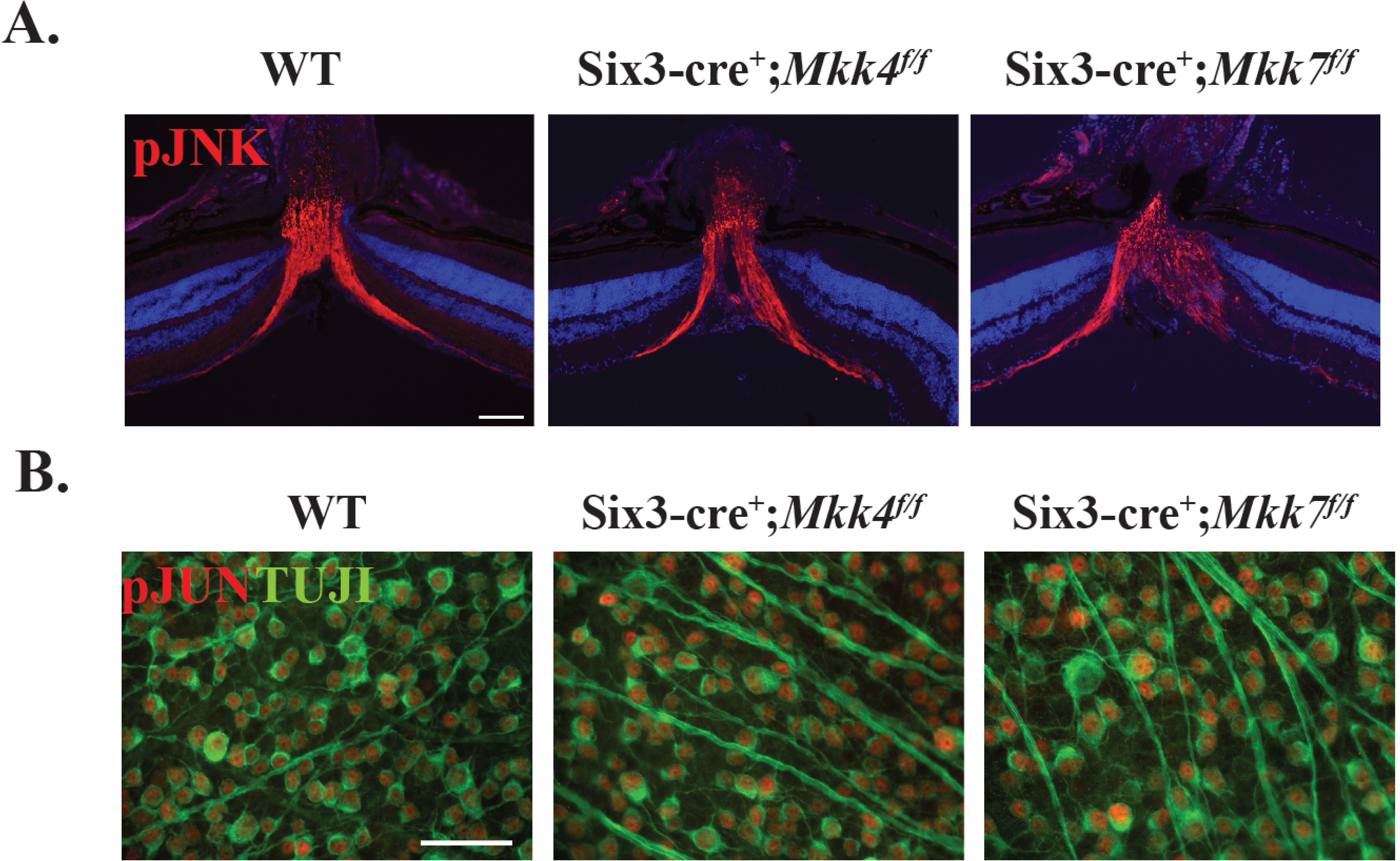
Deficiency of *Mkk4* or *Mkk7* does not prevent JNK-JUN signaling after axonal injury in RGC somas or axons. (**A**) Representative sections containing the optic nerve head demonstrate robust expression of pJNK two hours after CONC in WT controls and in *Mkk4* and *Mkk7* mutants (N≥3 per genotype). (**B**) Representative retinal flat mounts from each group reveal robust levels of pJUN in TUJ1 positive RGCs three days after CONC (N≥3 per genotype). Scale bars: 50μm.

### Deficiency of *Mkk4* or *Mkk7* protects RGCs after acute mechanical axonal injury

The MAPK signaling family has been previously shown to be an important pro-apoptotic signaling pathway after axon injury (e.g. ^6,7,38,39,40,41^). In order to determine if *Mkk4* and/or *Mkk7* is critical for axonal injury mediated RGC death, RGC survival was analyzed in mice deficient in *Mkk4* and *Mkk7* after CONC. *Mkk4* and *Mkk7* deficient retinas had significantly fewer dying RGCs (cleaved caspase 3 positive cells) five days after CONC as compared to wildtype controls even after correcting for the decreased number of RGCs in *Mkk4* and *Mkk7* deficient retinas (Fig. 4A,B). In addition, thirty-five days after CONC, *Mkk4* and *Mkk7* deficient retinas had a small, but significantly greater percentage of surviving RGCs as compared to wildtype controls (Fig. 4C,D; given as genotype, % survival ± SEM: WT, 16.0% ± 1%; *Mkk4* deficient, 51.5% ± 6.4%; *Mkk7* deficient, 30.4% ± 1.4%). However, it is important to note that this protection was not as robust as what was found in *Jun* deficient mice (~75%) at the same time point suggesting that both *Mkk4* and *Mkk7* can activate JNK-JUN dependent RGC death after axonal injury. This is consistent with the immunohistochemical results showing pJUN in RGCs after axonal injury in both *Mkk4* and *Mkk7* deficient mice (Fig. 3B).

**Figure 4.**
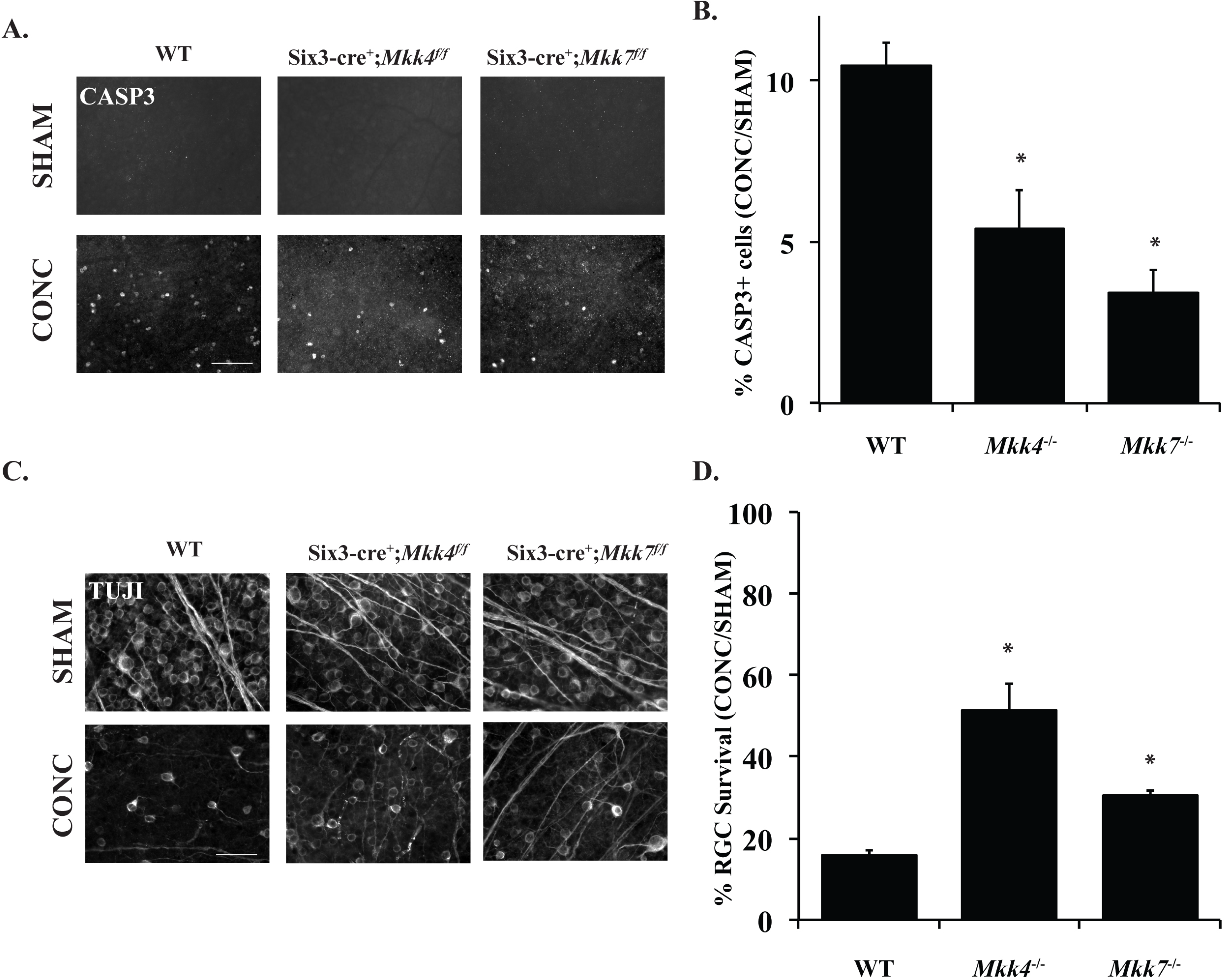
Deficiency of *Mkk4* or *Mkk7* provides moderate RGC protection after CONC. (**A**) Representative retinal flat mounts stained with cleaved caspase3 (CASP3) in eyes five days after CONC or sham surgery. (**B**) Significantly fewer apoptotic cells are present in *Mkk4* and *Mkk7* deficient animals as compared to WT controls following CONC. N≥4 per genotype per condition; * p ≤ 0.05. (**C**) Representative retinal flat mounts stained with TUJ1 following CONC or sham surgery. (**D**) Significantly more cells remain 35 days after CONC in *Mkk4* and *Mkk7* deficient retinas as compared to WT controls. N≥8 per genotype per genotype and condition. Scale bars: 50μm. Error bars represent S.E.M.

### Combined deficiency of *Mkk4* and *Mkk7* leads to severe alterations in retinal structure

Deficiency of *Mkk4* or *Mkk7* alone provided significant protection to RGCs after mechanical axon injury, however, neither deficiency by itself provided complete protection to RGCs. As *MKK4* and *MKK7* are the only molecules known to activate JNK, it appears that after mechanical axonal injury, MKK4 and MKK7 have redundant roles activating JNK. Moreover, as either deficiency alone did not cause severe retinal dysgenesis during development, removing both genes, and therefore any possible genetic redundancy, could allow for the role of MAP2Ks in axonal injury-induced RGC death to be tested. To test these possibilities, animals were generated that were deficient in both *Mkk4* and *Mkk7* in the retina (double conditional knockout using *Six3*cre). *Mkk4/Mkk7* deficient mice had an apparent complete lack of, or highly atrophic optic nerve (Fig. 5A), disrupted optic nerve head morphology (Fig. 5B), and a disorganized GCL layer (Fig. 5C; other retinal layers were also disorganized, discussed below). The lack of an optic nerve prevented analysis of axonal injury-induced death in *Mkk4/Mkk7* deficient mice. Surprisingly, despite the lack of an optic nerve, *Mkk4/Mkk7* deficient retinas still had TUJ1+ RGCs. However, there was clear evidence of abnormalities in RGC organization. Some areas of *Mkk4/Mkk7* deficient retinas displayed abnormal RGC axonal crossing, while other areas appeared relatively normal (Fig. 6A). An RGC clumping phenotype, more prevalent and severe than that seen in *Mkk7* deficient retinas, was observed as well (Fig. 6B).

**Figure 6.**
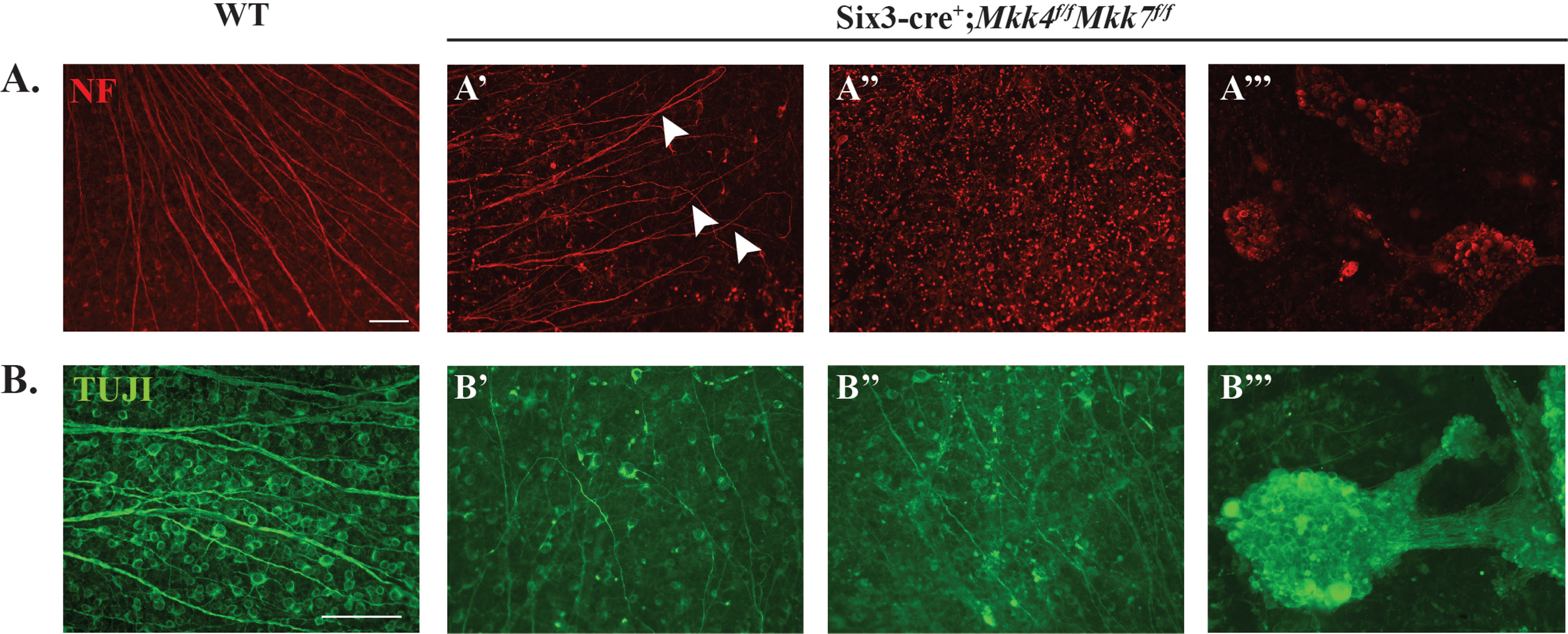
Fasciculation of RGCs and amacrine cells in retinas of *Mkk4/Mkk7* double deficient mice. Retinal flat mounts were stained with neurofilament (NF; **A**) and TUJ1 (**B**) to assess RGC organization. *Mkk4/Mkk7* double deficient mutants displayed phenotypic variability in RGC structure, with some areas exhibiting axonal overlap (arrowheads in **A’**) and irregular axonal trajectories (B’), others appearing relatively normal (**A’’**) and several areas consisting of RGC clumps (**A’’’, B’’’**). N≥4 per genotype; scale bars: 100μm.

**Figure 5.**
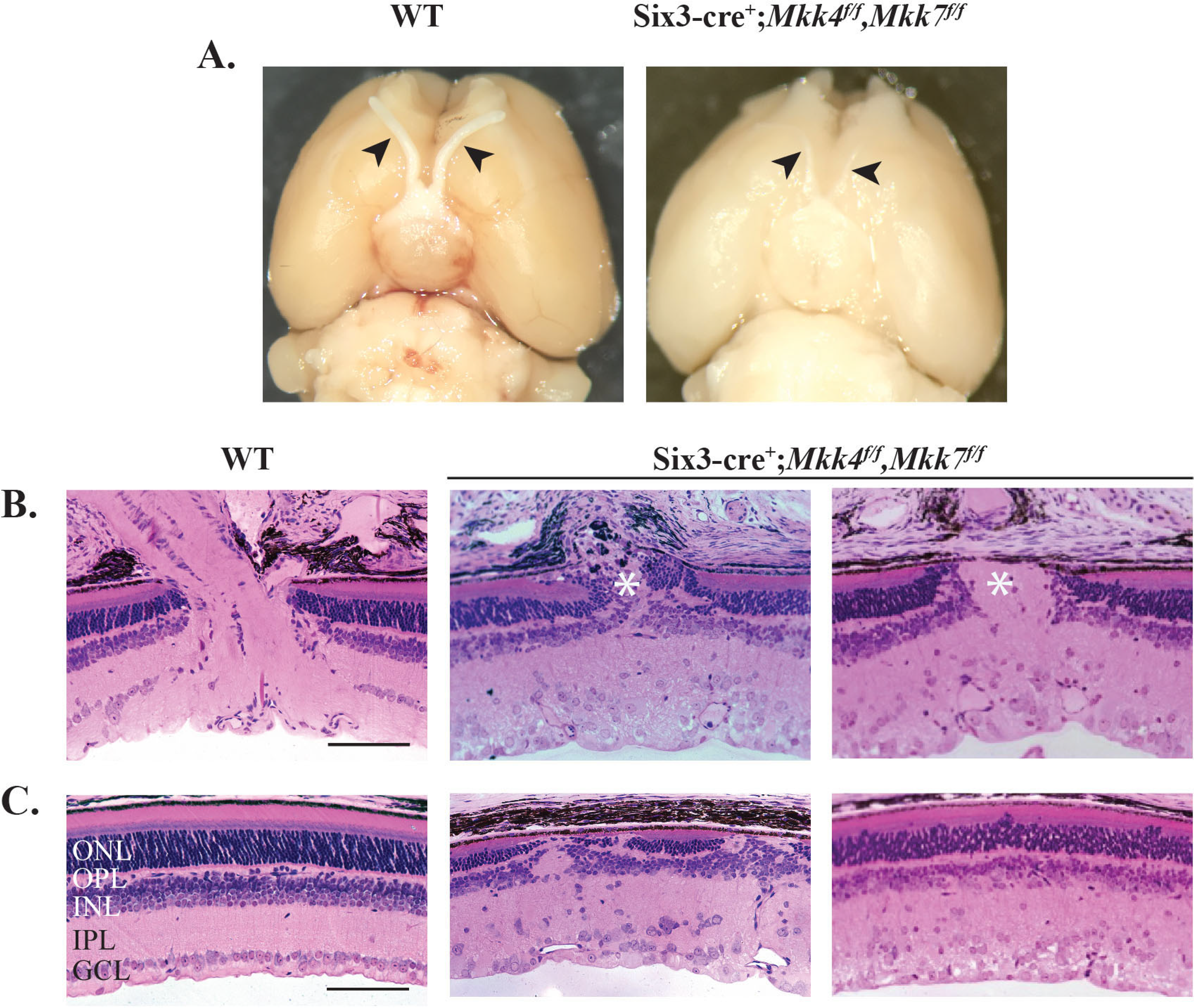
Combined deficiency of *Mkk4* and *Mkk7* causes failure of optic nerve formation and disruption of retinal lamination. (**A**) Brains were removed from adult WT and *Mkk4/Mkk7* double deficient mice with optic nerves intact (arrowheads). Near complete loss of both optic nerves was evident in all *Mkk4/Mkk7* mutant brains examined (N=8 per genotype). (**B**) Representative 3μM plastic sections stained with H&E further demonstrate severe optic nerve head abnormalities (asterisk) in *Mkk4/Mkk7* double deficient mice eyes (N=4 per genotype). (**C**) *Mkk4/Mkk7* double mutants display extensive disruption of all retinal layers (N=4 per genotype). Scale bars: 100μm.

Retinal sections also showed severe abnormalities in retinal lamination (Fig 5C). These abnormalities included areas of retinal photoreceptor thinning and areas of hyper- and hypoplasia in the inner nuclear layer. Overall, each layer of the retina appeared thinner compared to WT controls, and multiple cell types were misplaced. Interestingly, histological retinal phenotypes did not appear to worsen with age, as 9 month old *Mkk4/Mkk7* deficient mice had similar histological irregularities compared to the younger mice analyzed (Supp. Fig. 2). To further investigate the nature of the histological abnormalities, immunohistochemistry was used to identify retinal cell types and overall retinal organization. There were clear abnormalities in CHAT+ and calretinin+ amacrine cells. Particularly, these markers showed that the inner plexiform layer had areas of synaptic disruption (Fig. 7A). Horizontal cells labeled with calbindin-D-28K had a mild disruption of somal and dendritic organization (Fig. 7B). PKCα+ bipolar cells had abnormal somal organization in the inner nuclear layer, occasional bipolar cell nuclei present in the photoreceptor layer, and abnormal bipolar cell termination in the inner plexiform layer (Fig. 7C). SOX2+ Müller glia and amacrine cells were highly irregular in their somal distribution (Fig. 7D). Dopaminergic amacrine cell clumping and dendrite fasciculation was more severe in double MAP2K deficient retinas than in retinas deficient in only *Mkk4* or *Mkk7* alone (Fig. 7E), suggesting the involvement of compensatory mechanisms. Overall, there was a clear disruption of inner nuclear layer cell somal and synaptic organization.

**Figure 7.**
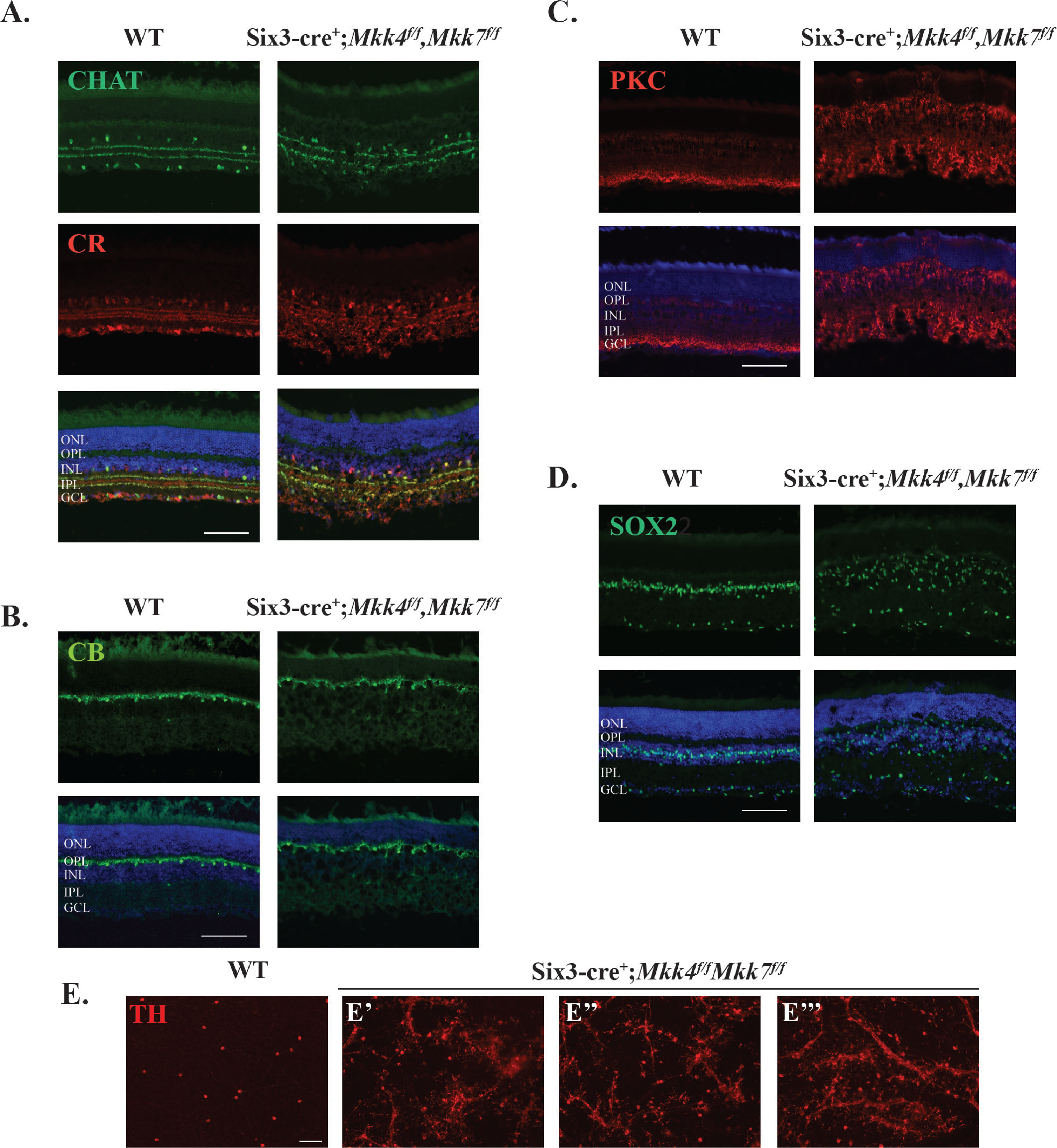
Combined deficiency of *Mkk4* and *Mkk7* leads to severe alterations in retinal structure. Representative sections of WT and *Mkk4/Mkk7* deficient retinas stained with specific cell markers show clear disorganization of amacrine cells (**A**), slight defects in horizontal cells (**B**), and abnormal distribution and morphology of bipolar cells (**C**) and Müller glia (**D**) in retinas of *Mkk4/Mkk7* double deficient mutants. Merged images with DAPI (blue) are shown below. N≥4 per genotype. (**E**) *Mkk4/Mkk7* double deficient flat mounts further display clumping and dendritic fasciculation of dopaminergic amacrine cells. N≥4 per genotype; scale bars: 100μm.

## Discussion

MAPK signaling is important for both retinal development and RGC apoptotic signaling after axonal injury. Herein, we demonstrate that MKK4 and MKK7 provide both overlapping and divergent roles in development and injury signaling in the retina. Single deficiency of *Mkk4* or *Mkk7* results in minor alterations of retinal lamination. RGC counts were significantly lower in adult *Mkk4* and *Mkk7* deficient retinas as compared to controls. After axonal injury, JNK signaling was not affected in *Mkk4* nor *Mkk7* deficient retinas, yet RGCs were mildly protected from apoptotic cell death in both mutants. However, RGC protection in the *Mkk4* and *Mkk7* single deficient animals was far from complete. To assess redundancy of MKK4 and MKK7 in both development and after injury, animals deficient in both *Mkk4* and *Mkk7* were developed. Together MKK4/MKK7 were found to play critical roles in the development and organization of all major retinal cell types. The severity of optic nerve dysgenesis combined with significant alterations in retinal organization in the *Mkk4/Mkk7* deficient retinas precluded examination of the contribution of both MKK4/MKK7 to pro-death signaling after axonal injury. Together these data suggest novel roles for both MKK4 and MKK7 in retinal development and in axonal injury signaling.

### Single *Mkk4* or *Mkk7* deficiency leads to multiple developmental abnormalities in the retina

Deficiency of either *Mkk4* or *Mkk7*, led to significant decreases in RGC number in adult retinas. Interestingly, this deficiency was not observed in retinas at P0. Throughout retinal development, an excessive number of cells are born that must undergo apoptotic pruning. Developmental RGC pruning occurs from P0 to P7, with a second, smaller wave of cell death occurring around P15^48,49,50^. Given the postnatal dropout of RGCs in *Mkk4* and *Mkk7* deficient animals, MKK4 and MKK7 likely contributes to RGC survival during the normal window of RGC death. Previously, the downstream effector molecules of MKK4 and MKK7, the JNKs, have been shown to regulate photoreceptor apoptosis during the final wave of cell death^14^. Thus, it is possible that MKK4 and MKK7 might play a similar role facilitating pruning of RGCs. Deficiencies in axonal transport or insufficient trophic support within the RGCs themselves may result in excessive loss of RGCs in the mature retina in *Mkk4* and *Mkk7* deficient mice^48,51^. Mild disruption of cell adhesion molecules may lead to the axonal fasciculations observed in the *Mkk7* deficient retinas. As axonal fasciculations were not observed in the *Mkk4* deficient retinas and *Mkk7* deficient retinas had significantly fewer RGCs, it is plausible that MKK7 plays a more significant role in axonal guidance and retinal pruning. Determining the overlapping and unique downstream effectors of MKK4 and MKK7 that contribute to retinal development will help define the required molecular cues for proper retinal lamination.

### Deficiency in both *Mkk4* and *Mkk7* causes severe retinal dysgenesis

In contrast to the *Mkk4* and *Mkk7* single deficient retinas, the *Mkk4/Mkk7* deficient retinas had severe alterations in retinal lamination and axonal projections. These data suggest that MKK4 and MKK7 have redundant roles in retinal development and that together these molecules play critical roles in retinogenesis. In addition to retinal lamination deficits, *Mkk4/Mkk7* deficient retinas had near absence of the optic nerve. Multiple downstream targets of MKK4/MKK7 activity may contribute to this dysmorphology. Previously, JNK signaling has been shown initiate bone morphogenetic protein-4 (BMP4) and sonic hedgehog (SHH) mediated control of paired-like homeobox transcription factor (PAX2), which contributes to closure of the optic fissure^12,52^. Netrin-1 also contributes to optic nerve head formation as deficiency of *netrin-1* leads to failure of RGC axons to exit the optic nerve leading to optic nerve hypoplasia^53^. Netrins serve as important axonal guidance cues in the nervous system and contribute to proper axon outgrowth and pathfinding^54^. JNK1 has been shown to be required for netrin-1 signaling and inhibition of JNK1 reduces netrin-1 dependent axonal projections and pathfinding in other neural systems^36^. Down syndrome cell adhesion molecule (DSCAM) is a netrin-1 receptor necessary for neurite arborization and prevention of abnormal neural fasciculations^55,56^. Axonal fasciculations and ectopic photoreceptor phenotypes similar to those in *Mkk4/Mkk7* deficient retinas were similarly observed in *dscam* deficient retinas^57^. Proper fasciculation is necessary for RGC axon pathfinding, however, as axons approach deeper targets in the brain, they must disassociate from their neighbors^58^. It is possible that in *Mkk4/Mkk7* mutants, the extrinsic cues directing axon divergence are disrupted, resulting in abnormal fasciculation in the retina. *Jnk2/3* deficient mice and *Jun* deficient mice do not have aberrant lamination or optic nerve head dysgenesis, which further supports the idea that morphologic differences observed in the *Mkk4/Mkk7* deficient mice are likely due to JNK1, or JUN independent signaling mechanisms^7,59^. Future study to evaluate the downstream signaling mechanisms of MKK4 and MKK7 will likely reveal key contributors to retinal and optic nerve development. Despite the lack of optic nerve formation in the *Mkk4/Mkk7* deficient animals, RGCs survive within the retina. Normal pathfinding and neurotropic support is essential for RGC survival, thus RGC survival in the absence of optic nerve formation likely is due to the multiple roles played by MAPK signaling^14,60,61^. JNK and JUN contribute to both pro-survival and pro-apoptotic signaling^21,62,63,64,65^. RGC survival in the *Mkk4/Mkk7* deficient retinas despite impaired optic nerve formation might therefore be due to decreased pro-apoptotic signaling.

### MKK4 and MKK7 are both involved in axonal injury-induced RGC death

Pro-apoptotic MAPK signaling is also an important component of molecular signaling after axonal injury in RGCs (e.g. ^6,7,8,21,22,23,24,39,40,41^). Similar to other MAPK family members, single deficiency of *Mkk4* or *Mkk7* provides significant protection to RGCs after axonal injury. The level of protection observed, however, does not phenocopy deficiency of other MAPK signaling molecules. Deficiency of the upstream MAP3K, *Dlk*, appears to provide greater protection to RGCs after optic nerve crush than that observed in this study^6,39,40,66^. Deficiency of downstream targets such as combined deficiency of *Jnk2/3* and deficiency of *Jun* also appears to afford greater protection to RGCs than *Mkk4* or *Mkk7* deficiency alone^7,59^. Together these data suggest that MKK4 and MKK7 likely have redundant roles in pro-apoptotic signaling after axonal injury.

While both MKK4 and MKK7 are known to activate the JNKs, MKK4 additionally regulates p38 ^4,67^. Activation of p38 also occurs after axonal injury in both neurons and glia^11,47,68,69,70^. Inhibition of p38 signaling has previously been shown to provide mild protection to RGC somas and reduce axonal transport deficits following axonal injury^11,68,69^. Activation of the p38 arm of MAPK signaling can also directly activate *Ddit3* (DNA damage inducible transcript 3) which encodes the protein CCAAT/enhancer binding homologous protein (CHOP)/GADD153, a key mediator of endoplasmic reticulum stress^71,72^. DDIT3 has been independently shown to be important for pro-apoptotic signaling after axonal injury^59,73,74^. In order to understand the molecular signaling cascade contributing to RGC death after axonal injury, it will be necessary to parse apart the downstream signaling contributions of MKK4 and MKK7.

Future studies should also evaluate combined *Mkk4/Mkk7* deficiency in a temporally controlled conditional knockout animal (allowing for normal retinal and optic nerve development) to determine if deficiency of these molecules together might provide greater protection to RGCs after injury than either alone. It is tempting to consider that dual deficiency of *Mkk4/Mkk7* may be more protective of RGCs than deficiency of other downstream targets alone, as combined deficiency of *Mkk4/Mkk7* may alter multiple molecular signaling pathways that have been previously shown to be important for RGC death including JNK, p38, and endoplasmic reticulum stress signaling^7,11,59,75^.

Future investigation of the unique and overlapping roles of MKK4 and MKK7 must consider the downstream targets of their activation. MKK4 and MKK7 are known to preferentially target different amino acid residues on the JNKs. MKK4 targets the tyrosine residue while MKK7 preferentially phosphorylates the threonine residue^76,77^. Analysis of the transcriptome of MKK4 and MKK7 is necessary to identify specific downstream targets. Such inquiry would determine if the downstream effectors leading to developmental phenotypes in deficient animals are perhaps independent from those downstream effectors mediating pro-apoptotic signaling after optic nerve crush. Furthermore, it would identify the key molecular signaling pathways that contribute to RGC death after axonal injury.

## Conclusion

MKK4 and MKK7 are members of the MAPK pathway important for both retinal development and the injury response following axonal insult. While we have shown that MKK4 and MKK7 are required for maintaining RGC survival, future studies should examine the cellular and molecular mechanisms underlying the observed postnatal dropout in single deficient animals. Single deficiency of *Mkk4* and *Mkk7* offer mild protection for RGCs against optic nerve injury, though there was likely genetic compensation allowing for pro-apoptotic signaling to proceed. Due to the severity of developmental defects in the double deficient mice, possible additive protection following mechanical insult was not assessed. Additional experiments employing an inducible double *Mkk4/Mkk7* knockout would allow for proper optic nerve formation and subsequent analysis of RGC survival following injury. This would define the aggregate role of these MAP2K molecules in a glaucomatous-relevant injury response and determine the specificity of the cellular response in both development and following axonal injury.

## Acknowledgements

The authors would like to thank Drs. C. Tournier and R.J. Davis for generously providing the *Mkk4* and *Mkk7* floxed alleles. This work was supported by EY018606 (RTL), EY026301 (RLR), EY007125 (RLR) and Research to Prevent Blindness, an unrestricted grant to the Department of Ophthalmology at the University of Rochester Medical Center. The funding agencies had no role in the design of the study and collection, analysis, and interpretation of data and in writing the manuscript.

## Supplementary Information

**Supplementary Figure S1. Validation of *Mkk4* and *Mkk7* deletion using *Six3*cre.**

Representative western blots showing levels of MKK4 and MKK7 protein in respective *Six3*cre driven conditional knockouts. There was an absence of MKK4 protein and a clear reduction of MKK7 protein compared to littermate WT controls (N≥4 per genotype).

**Supplementary Figure S2. Retinal lamination in aged *Mkk4/Mkk7* double mutants.**

Representative 3μM plastic sections stained with H&E reveal additional retinal deterioration does not occur over time in *Mkk4/Mkk7* deficient mutants aged to 9 months (N=4 per genotype). Extensive disruption of all retinal layers is similar to that observed in *Mkk4/Mkk7* deficient mutants examined at 2 months of age (refer to Fig. 5).

